# Glycolytic repression and reduced GLP-1 secretion by active Farnesoid X Receptor in enteroendocrine L cells achieved via PKLR

**DOI:** 10.1101/242313

**Authors:** Kristoffer Niss, Søren Brunak

## Abstract

Rise in intestinal glucose increases GLP-1 secretion by enteroendocrine L cells. GLP-1, in turn, stimulates insulin secretion. Farnesoid X Receptor (FXR) represses this pathway by manipulating the L cell glycolysis, thus reducing insulin- and GLP-1 secretion. The mechanism by which FXR manipulates the L cell glycolysis is unclear. In this study, we construct an L cell specific protein-protein interactome and identify all significantly active protein complexes, inferred by co-expression scores, in FXR-activated and control L cells. Contrary to previous reports, we find extensive glycolytic enzyme activity in FXR-activated L cells. We present how FXR’s repression of the glycolytic enzyme, pyruvate kinase (PKLR), is causing the reduction in glycolytic activity. This mechanistic insight may aid development of drugs targeting the GLP-1 pathway.

## Introduction

Enteroendocrine L cells are chemosensors situated in the epithelium of the human intestine, characterized mainly by their ability to release the hormones glucagon-like peptides 1 and 2 (GLP-1 and −2) ^1,2^, and additionally, peptide YY and insulin-like peptide 5 ^2,3^. L cells have been shown to sense bile acids ^4^, monosaccharaides ^1^, fatty acids ^5^ and possibly microbial metabolic products ^6^. L cell hormones are secreted in response to changes in the intestinal microenvironment. GLP-1 secretion specifically has been linked to increased concentrations of glucose ^1^ and long chain fatty acids ^5^. Multiple organs are affected by the pleiotropic effects of GLP-1 when secreted ^7,8^. Increased glucose uptake and appetite regulation has been shown, but of utmost importance is GLP-1’s incretin effect, in which insulin secretion is amplified by the activation of GLP-1 receptors on pancreatic β cells ^7,8^. A reduction in the strength of the incretin effect has pathological implications, as seen in type 2 diabetes (T2D), where a decrease in meal-induced GLP-1 secretion can, in part, explain the disease hallmark of insufficient insulin secretion ^8,9^.

The mechanism by which L cells intracellularly translate increased glucose concentration (glucose-sensing) into GLP-1 secretion is not yet fully understood. But with the growing interest in GLP-1 pharmacotherapy ^7,10^, the mechanism is of considerable interest. In T2D, for an example, the incretin effect and glucose-lowering property of GLP-1 are used to increase insulin secretion and normalize hyperglycemia^7^. This is most commonly done using GLP-1 receptor agonists that target pancreatic β cells and stimulate insulin production ^7,10^. However, by delineating the mechanism that links glucose-sensing and GLP-1 secretion, innovative approaches may be discovered. In this regard, Trabelsi et al. recently demonstrated how activated farnesoid X receptor (FXR) represses L cell GLP-1 secretion *in vivo* by reducing glycolytic ATP production ^11^. FXR is activated by bile acids ^11,12^, and it was furthermore presented how the sequestration of bile acids could prevent FXR activation, consequently improving GLP-1 secretion. Trabelsi et al. thus showed how one can manipulate pathways upstream of GLP-1 secretion and hereby evoke raised endogenous GLP-1 secretion overall.

In this study, we want to identify key proteins in L cells that link glucose-sensing to GLP-1 secretion. Protein products of most genes carry out their function in concert through physical interactions ^14^. We therefore use the physical interaction neighbourhood of proteins (in terms of millions of experimentally derived pairwise interactions) to add context to the gene expression changes we observe in L cells, and thus filter noise ^15^. For the latter, we use the publicly available GLUTag ^13^ L cell line microarray gene expression data from Trabelsi et al. ^11^. It is a dual-condition data set, where FXR is either activated (FXR-activated) or not (control). Genes with expression differences between the conditions are possible targets of FXR, and since active FXR represses GLP-1 secretion via the glycolysis, those genes may link glucose-sensing to GLP-1 secretion.

By integrating L cell gene expression data with a recently published human physical protein-protein interactome, InWeb_IM ^16^, we first create an L cell-specific interactome and subsequently list and rank its protein complexes. Such complexes are strong predictors of functional association between proteins ^17^. It is advantageous to work with protein complexes rather than hand-curated pathways, as complexes can be overlapping ^18^ and cell- or tissue specific ^19^, thus depicting the flexibility of interacting proteins more accurately. Moreover, protein interactions are often derived experimentally in large screens, making it possible to incorporate entirely novel components in already described pathways. The relative expression of the protein units in a complex, its stoichiometry, can then be used to assess the activity of each protein complex. High gene co-expression, which infers a balanced protein complex stoichiometry, is an indicator of activity or importance ^19–23^. Using a rigorous and untargeted selection scheme, we identify the most robustly co-expressed protein complexes and thus those that are the most active in a specific context. Disruption or gain in activity between FXR-activated- and control L cells consequently map protein complexes that are affected by FXR activation.

We reveal that active FXR is not repressing the glycolysis as a whole, since the glycolysis show high activity in FXR-activated L cells. Instead, we present how active FXR exclusively targets the glycolytic enzyme liver-type pyruvate kinase (PKLR), hereby provoking decreased glycolytic activity and reduced GLP-1 secretion.

## Methods

### Microarray GLUTag L cell line gene expression data

The expression data used, stem from the Trablsi et al. study ^11^ and was downloaded from ArrayExpress (id: E-MTAB-2199). It consists of four control L cell samples and four GW4064-treated L cell samples. GW4064 is a FXR agonist and activator. Preprocessing was done using the R package *Limma* ^24^ by following the *Limma* manual’s Agilent microarray example. ENMUST annotations were mapped to human UniProt annotations using the functions *gorth* and *gconvert* from gProfileR ^25^.

### The L cell specific interactome and repertoire of protein complexes

We applied the method described in Pedersen et al. ^19^ to construct the protein interactome and protein complex repertoire. To make the complexes, two complementary clustering methods were used: spoke complex construction ^20^ and clusterONE ^18^, which produced 3,356 spoke complexes and 505 dense clusterONE complexes. As a modification to the Pedersen et al. workflow, we applied the complex size restrictions of > = 3 and < = 25 proteins, as this range better reflects the size distribution of mammalian protein complexes (See **Figure S2**). Finally, we reduced the protein complexes to a set of 1,221 after size limits were applied and highly overlapping complexes were merged.

### Protein complex co-expression score

To calculate the co-expression of two arbitrary proteins, we used an approach described previously ^19–22,26^. If proteins *x* and *y* were present in the same protein complex and had a physical interaction according to InWeb_IM, we calculated the gene expression correlation between protein *x* and *y* using the Pearson’s correlation coefficient (PCC_xy_):

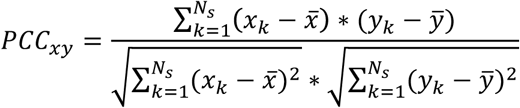
where *N_s_* is the number of samples and *x_k_* is the expression value of protein *x* in sample k. Similarly, *y_k_* is the expression value of protein *y* in sample *k*. For each protein complex, we then aggregate the pairwise PCC values into a *co-expression score* using a weighted average (see supplementary materials). We weigh each PCC value on the confidence score of the protein-protein interaction and the signal-to-noise ratio of the correlation (see the section below regarding the signal-to-noise ratio and supplementary materials regarding the *co-expression score)*.

### The signal-to-noise ratio

Gene expression correlation is a quantification of how similar the between-sample variation of two genes are. We assess, if a pair of genes is highly- and lowly expressed in the same samples. However, the between-sample variation that we observe is not only due to biological effects, but also due to measurement noise ^27^. To differentiate between correlations based on mainly measurement noise and correlations based on biological effects, we calculated a signal-to-noise ratio (SNR) for each correlation. The majority of gene expression fluctuation is due to measurement noise, and the noise level is dependent on each genes’ mean expression level ^27^. We modeled the measurement noise of our expression data using a simple linear regression model, calculating the square root of the standard deviation (SD) dependent on the average gene expression (**Figure S3**, black line):

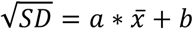
where 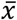 is the average gene expression. We found: a = −0.01149 and b = 0.52252. The estimated measurement noise of gene *k* (*noise_k_*) can then be calculated as:

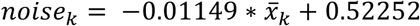
where 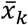 is the average expression value for gene *k*. The amount of biological signal for gene *k* (*signal_k_*) can then be calculated as:

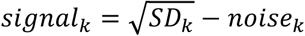
where *SD_k_* is the standard deviation of gene *k*. If *signal_k_* is less than zero, we set it to zero before calculating the SNR of gene *k:*

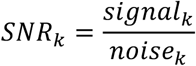

The SNR of the correlation between gene *k* and gene 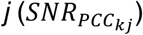 is:

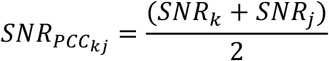

To calculate the SNR for a whole protein complex (*SNR_complex_*):

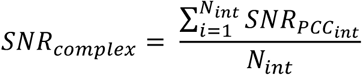
where *N_int_* is the number of interactions and 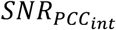 is the SNR for a correlation between two proteins that have a physical interaction (*int*) in the complex.

### Hierarchical clustering

Clustering of the protein complexes was done by first calculating the average shortest path between the proteins of all complex-pairs using the *networkx* Python package ^28^, and then hierarchically cluster the complexes using the R function *hclust*.

### Permutation test: Significant complex co-expression score relative to random protein complexes

For each protein complex, a permutation test asked, if co-expression scores higher than or equal to the observed L cell co-expression score could occur in a random background of 10,000 co-expression scores. Random co-expression scores were calculated by assigning random expression values from our expression data to the complex in question, and recalculating the co-expression score. The number of times a random co-expression score was higher than or equal to the observed score divided by 10,000 was then assigned as the p-value. Finally, multiple-testing p-value adjustment of Benjamin-Hochberg 5% False Discovery Rate was applied.

### Permutation test: Significant co-expression between the glycolytic enzymes and PKLR relative to a random protein

The following was done for each L cell state. The average correlation between PKLR and the other glycolytic enzymes in L cells was calculated using Pearson’s correlation coefficient (PCC). A permutation test was then used to ask, if scores higher than or equal to the observed average correlation could occur in a random background of 100,000 average correlations. Random average correlations were calculated by assigning random expression values to PKLR for each calculation, keeping the observed values for each glycolytic enzyme, calculating the PCC and finally taking the average. The number of times a random average correlation was higher than or equal to the observed value divided by 100,000 was then assigned as p-value.

### Gene ontology enrichment analysis

Gene ontology (GO) analysis was performed using the gProfileR Bioconductor R package ^25^. The proteins of the L cell interactome were used as custom background, a strong hierarchical filtering was enforced and only the enrichment of GO biological processes was calculated.

### Differential expression analysis

We used the *Limma* framework ^24^ for normalization (quantile, scale, cyclicloess and none) and as one of the chosen approaches of differential expression analysis. As the second differential expression approach, we used the build-in R function for the Welch Two Sample t-test.

## Results

### L cell-specific interactome and protein complexes

To construct a repertoire of L cell-specific protein complexes, we first created an L cell-specific protein interactome by removing all proteins that were not expressed in the GLUTag L cell microarray data from the protein-protein interactome InWeb_IM ^16^. We hereafter computationally decomposed the L cell interactome into 1,221 overlapping protein complexes using a modified version of the method described by *Pedersen et al*. ^19^, where the complex size limits were changed (>= 3 and <= 25) to better reflect the distribution of mammalian protein complexes (**Figure S2**).

After protein complex repertoire construction, we calculated complex-by-complex co-expression scores, which was done separately for FXR-activated and control L cells. For each physical protein-protein interaction in a complex, we obtained the protein pair co-expression by calculating the absolute Pearson’s correlation coefficient (PCC) using the expression values of the genes that encode the proteins. Subsequently, we aggregated the absolute PCC values of each protein complex into a complex-specific *co-expression score* by calculating the weighted-average of the absolute PCC values.

### Untargeted selection of active protein complexes

We searched for the presence of protein complexes with very high co-expression scores, i.e. activity, in FXR-activated or control L cells. Initially, we visualized the co-expression score distributions of the two conditions (**Figure 1A-B**). This showed that both conditions had similarly shaped distributions with considerable right tails (**Figure 1A-B**, asterisks). Moreover, the tails contained protein complexes with co-expression scores that were exceedingly higher than the random background co-expression level (**Figure 1**, black distributions).

**Figure 1.**
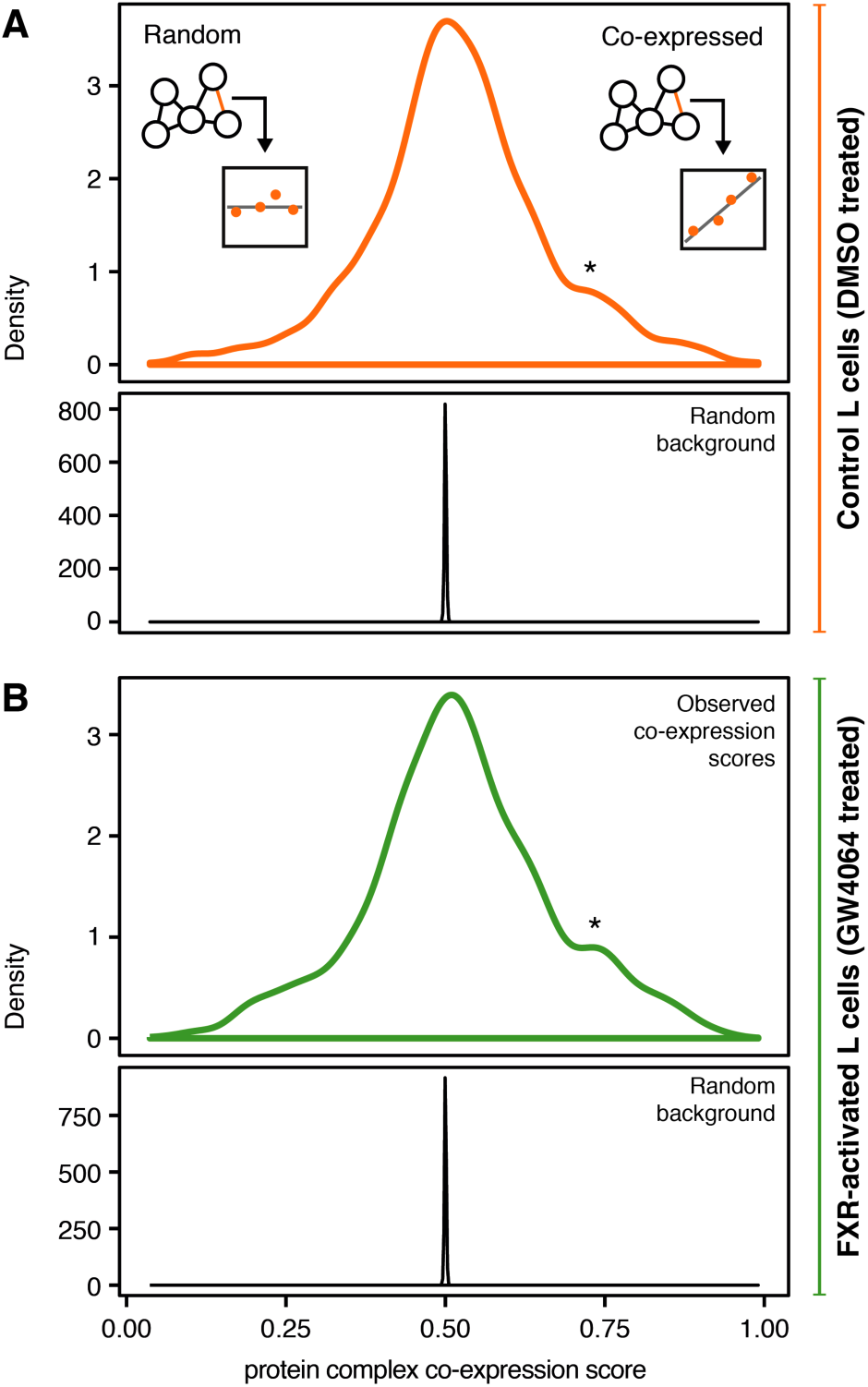
Protein complexes with high co-expression present in both L cell states. The density distribution of all protein complex co-expression scores in control (DMSO-treated) L cells is seen in **A** and for FXR-activated (GW4064-treated) L cells in **B**. Asterisks (*) indicate distribution shoulders containing highly co-expressed protein complexes, while the black distributions located in the bottom part of **A** and **B** are background distributions based on permutated expression values.

We therefore tested the statistical significance of all protein complex co-expression scores in both conditions, using a permutation test (10,000 iterations, applying 5% FDR p-value adjustment). This revealed that 7 and 14 complexes showed significant co-expression scores in, respectively, control L cells (**Figure 2B**, red bars) and FXR-activated L cells (**Figure 2D**, red bars). To ensure that the significantly co-expressed protein complexes in each condition were concurrently based on strong gene expression variation, we calculated the average signal-to-noise ratio (SNR) of the PCC values in each complex, hence termed *complex SNR* (**Figure 2C and 2E**). Having a high protein complex SNR signifies that the complex primarily contains PCC values that have been calculated using above-noise between-sample variation. Complexes with high SNR hereby have a higher likelihood of being controlled by a strong biological effect.

**Figure 2.**
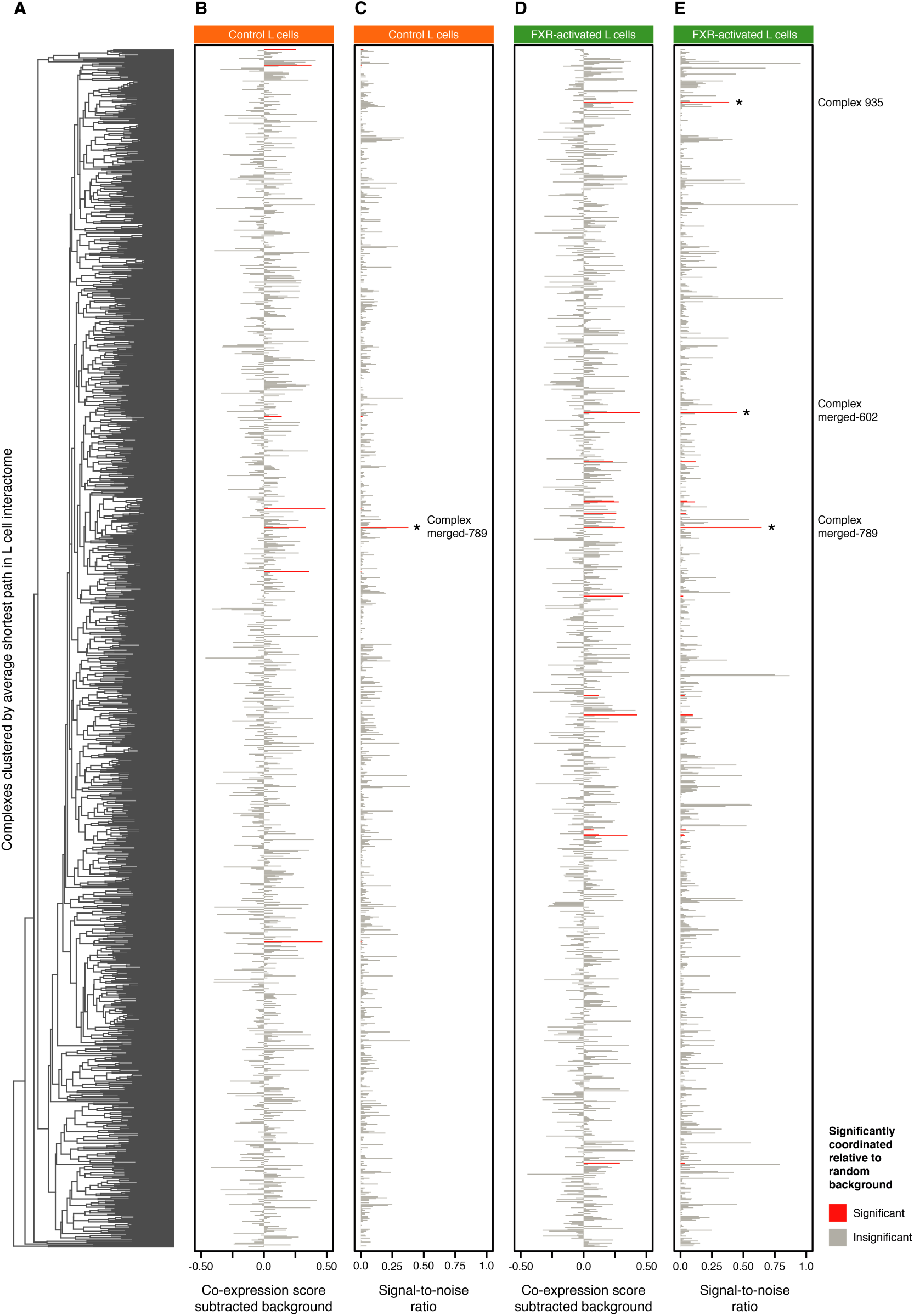
Overview: Three protein complexes are singled out as highly active. By displaying the co-expression scores and the signal-to-noise ratios (SNR) of each protein complex, complex “merged-789”, “merged-602” and −935 stood out (asterisks). The protein complexes are ordered vertically in **A** using a hierarchical clustering dendrogram based on the average shortest path between the complexes in the L cell interactome. The co-expression score subtracted the background co-expression level for each complex is shown for control L cells in **B** and for FXR-activated L cells in **D**. Grey bars indicate insignificant co-expression scores, while red bars indicate significant co-expression scores (perm. test, 5% FDR) in all panels. The signal-to-noise ratio (SNR) for each co-expression score is shown for control L cells in **C** and for FXR-activated L cells in **E**. Asterisks (*) indicate complexes with both a significant co-expression score and a notably high SNR.

**Figure 2** is a comprehensive overview of the co-expression scores and SNR’s for all protein complexes in both conditions. From this overview, it was clear that complex “merged-789” stood particularly out in control L cells (**Figure 2C**, asterisk). Complex “merged-602”, complex 935 and, appearing in both conditions, complex “merged-789” stood out in FXR-activated L cells (**Figure 2E**, asterisks). These protein complexes both had significant co-expression scores and particularly high SNRs, inferring activity. Network visualizations and gene ontology analyses of complex “merged-602” and −935 can be seen in **Figure S4**.

### Complex “merged-789” depicts an active glycolysis in FXR-activated L cells, supported by DE analysis

Complex “merged-789” arose from our analysis as the most intriguing protein complex. It had, as the only complex, a significant co-expression score in both states (control p-value 0.0E-4; FXR-activated p-value 0.0E-4), while also having the highest SNR of all protein complexes in both states (control SNR 0.38; FXR-activated SNR 0.65). The proteins of complex merged-789 are back-to-back glycolytic enzymes, converting glyceraldehyde 3-phoshate to phosphoenolpyruvate in four steps, i.e. steps 6-9 of the classical glycolysis pathway (**Figure 3A**). Consequently, the protein complex is significantly enriched for the gene ontology term canonical glycolysis (GO:0061621, p-value 4.18E-12).

**Figure 3.**
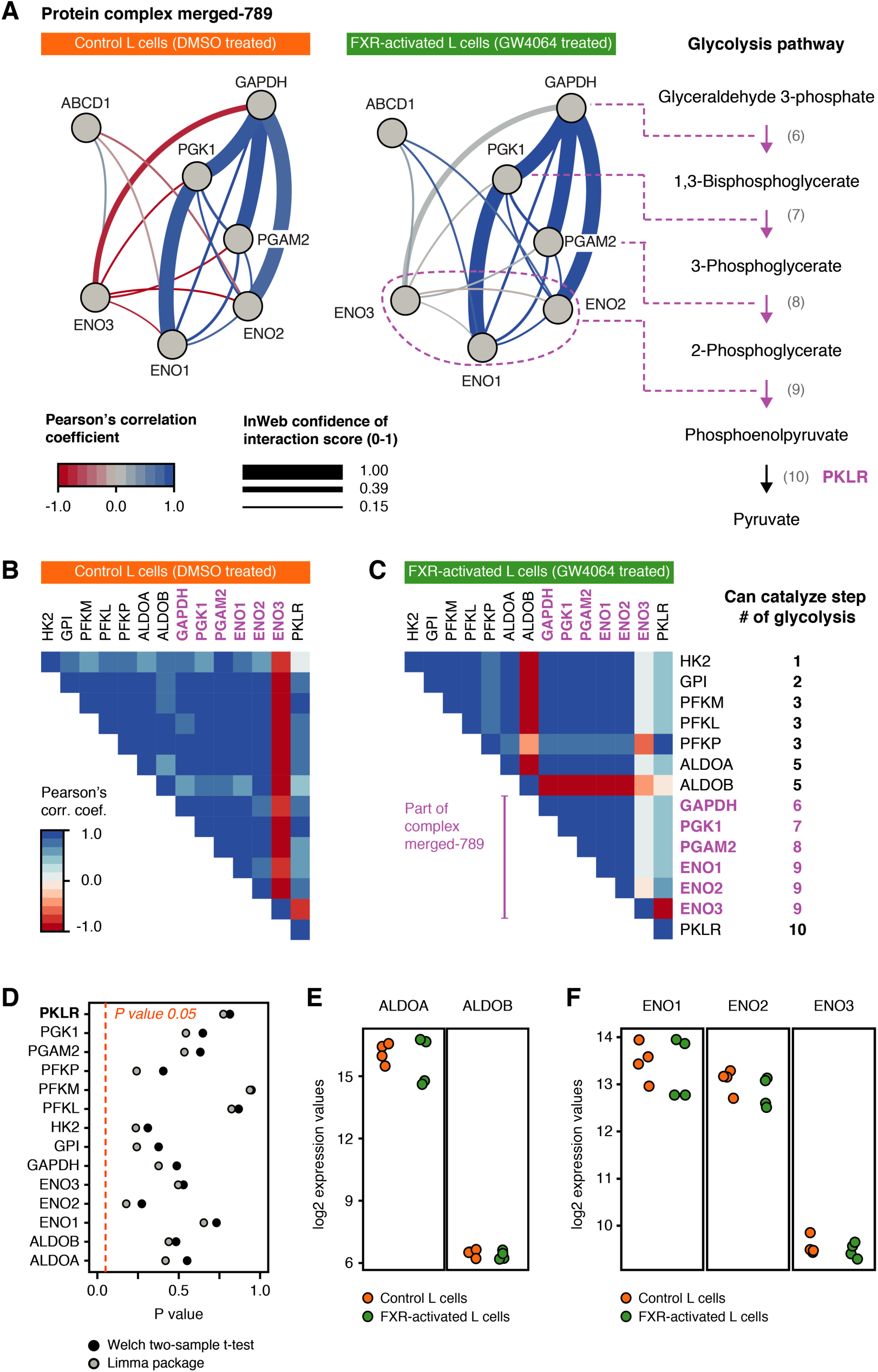
Complex merged-789 reveals an active glycolysis in FXR-activated L cells. Network visualizations of protein complex “merged-789” are shown in **A** for control L cells (orange) and FXR-activated L cells (green). Edges between the proteins are experimentally validated physical PPI’s from InWeb_IM, where edge thickness indicates the interaction confidence score and edge color indicates expression correlation (Pearson). The pathway representation (**A**, right) highlights the central position of complex “merged-789” in the glycolysis. Correlation matrices (Pearson) of all glycolytic enzymes are visualized for control L cells (**B**) and FXR-activated L cells (**C**). Blue indicates high co-expression, while red indicates negative co-expression. Most of the glycolytic enzymes display correlation values of 0.9 or higher (blue). The proteins of complex “merged-789” are in purple. Plot **D** illustrates the absence of significantly differentially expressed glycolytic protein. None of the 14 glycolysis-related proteins has an unadjusted p-value below 0.05 independent of statistical test (*limma* and Welch two-sample t test). See **Figure S5** for details. Expression strength plots of ALDOA and ALDOB (**E**), and ENO 1-3 (**F**) are also shown, where the color indicates the L cell state (orange: control, green: FXR-activated).

Glycolysis is a centerpiece of L cell biology, and FXR specifically has been shown to reduce glycolytic ATP production in L cells by Trabelsi et al., which was linked to reduced GLP-1 secretion ^1,8,11^. Trabelsi et al. furthermore proposed that the underlying mechanism of this effect is a transcriptional decrease of multiple glycolytic enzymes. Yet, evidence of this mechanism is not observed in our analysis. Instead, we find the glycolytic enzymes of protein complex “merged-789” meticulously co-expressed (**Figure 3A**, PCC > 0.9). If we broaden the view to encompass the complete glycolysis (**Figure 3B-C**), we find almost every glycolytic enzyme co-expressed (PCC >= 0.9). These observations indicate an active pathway.

Investigating this contradiction, we thoroughly redid the differential expression (DE) analysis between FXR-activated and control L cells that prompted Trabelsi et al. to make their “glycolytic repression” proposal. We performed the DE analysis using sixteen different strategies, which were permutations of: two gene subsets, two variants of statistical tests and four types of between-sample normalizations (**Figure S5**). We found no significantly differentially expressed genes (p-value < 0.05) related to the GO term “canonical glycolysis”, independent of applied strategy (**Figure 3D** and **S5A-H**). We therefore find evidence of an, on the whole, transcriptionally active glycolysis in FXR-activated L cells.

### PKLR is affected by the activation of FXR and is likely causing the glycolytic repression

However, as the decrease in glycolytic ATP production following FXR activation is still evident, a mechanism must link the glycolysis to FXR. In FXR-activated L cells, three glycolytic enzymes show deviating expression patterns from the overall glycolytic co-expression pattern: ALDOB, ENO3 and PKLR (**Figure 3C**). Reduction in co-expression is a predictor of reduced functional activity ^19–23^, which may cause the glycolytic repression. However, the reduced co-expression of ALDOB and ENO3 likely has no functional impact, as correspondingly ALDOA and ENO1-2, can catalyze the same glycolytic steps (**Figure 3C, right**). Furthermore, the subordinate roles of ALDOB and ENO3 is underlined by their low expression levels (**Figure 3E-F**). PKLR, on the contrary, single-handedly catalyze step 10 of the glycolysis (**Figure 3C, right**). This makes PKLR an essential glycolytic enzyme, and its deviating expression pattern may have a functional impact.

In **Figure 4A**, we have visualized the expression patterns of the glycolytic enzymes in FXR-activated L cells. Here, PKLR shows a deviating pattern (**Figure 4A, red line**). By calculating the average correlation (avg. PCC) between PKLR and the other glycolytic enzymes in both FXR-activated- and control L cells, and hereafter quantifying the difference, we find a 42% decrease in correlation (Avg. PCC: control=0.57 and FXR-activated=0.33). Thus, PKLR becomes less co-expressed with the glycolysis after FXR is activated. **Figure 4B** shows the difference in correlation between PKLR and each glycolytic enzyme in FXR-activated- and control L cells as a bar plot, while **Figure 4C** shows the correlations in both conditions as a heat map.

**Figure 4.**
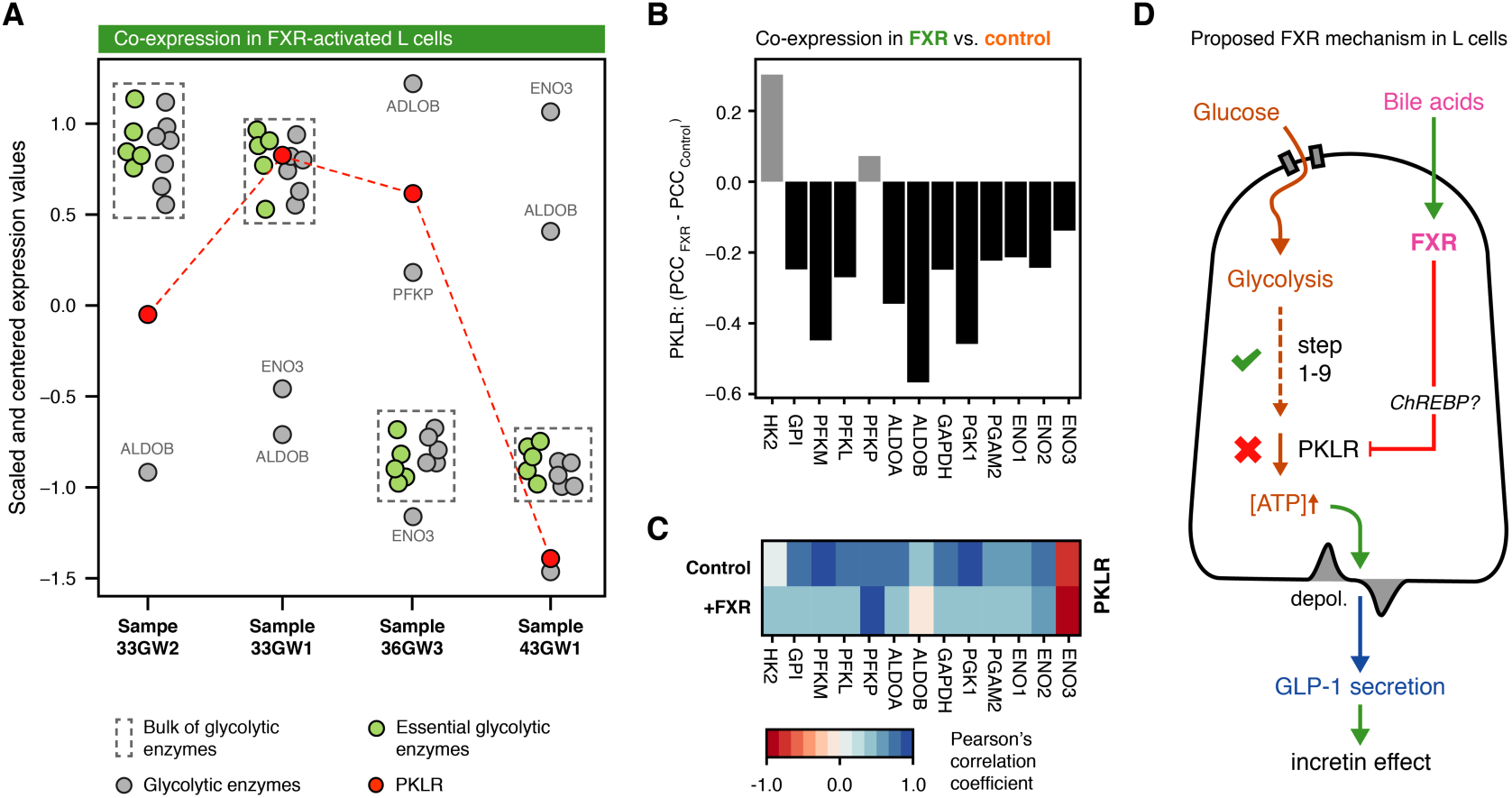
PKLR is the target of activated FXR. Scaled expression values of the glycolytic enzymes in FXR-activated L cells are shown in **A**. The x-axis shows each sample and the y-axis shows the expression. The essential glycolytic enzymes (green) are highly co-expressed, while PKLR (red) and the non-essential glycolytic enzymes (grey) are uncoordinated. A barplot (**B**) indicates the change in correlation between PKLR and all other glycolytic enzymes in control L cells relative to FXR-activated L cells. When FXR is activated, 11/13 proteins show a decrease in correlation to PKLR. A heatmap (**C**) shows the correlation between PKLR and the other glycolytic enzymes in each L cell state, as indicated by color. The schematic (**D**) illustrate our proposed mechanism of FXR in L cells: FXR is activated by bile acids and reduces glycolytic ATP output by targeting PKLR expression, possibly via the transcription factor ChREBP. This causes a decrease in GLP-1 secretion, “incretin” effect and, finally, insulin stimulation.

We furthermore asked, if the average correlation between PKLR and the other glycolytic enzymes was significant in FXR-activated or control L cells (permutation test, 100,000 iterations). The average PKLR correlation turned out to be very significant in control L cells (p-value 0.00011), while a lower confidence was seen in FXR-activated L cells (p-value 0.01810). We believe that these findings are indications of an FXR-driven disruption of PKLR’s gene expression. As PKLR has a key bottleneck position, where the regulation of PKLR can regulate the glycolysis globally, the reduction in glycolytic ATP production may be due to PKLR gene expression regulation exclusively. We have formulated our hypothesized regulation of the glycolysis by active FXR via PKLR in a schematic in **Figure 4D**.

Our PKLR hypothesis is further strengthened due to the fact that an identical mechanism has been observed in another cell line. A study by *Caron et al*. ^29^ showed that active FXR physically blocks glucose-induced transcription factor ChREBP in immortalized human hepatocytes (IHH), such that ChREBP in turn cannot induce the expression of PKLR specifically, when the concentration of glucose rises. The blocked PKLR expression hereby repressed glycolysis as a whole. Their observations in IHH are in alignment with our findings in L cells. Given the high conservation of protein function, even between distant species ^30^, we find it probable that the same protein, regulating the same pathway, does this by a similar mechanism, even though the cell types in the two studies are different.

## Discussion

Trabelsi et al. ^11^ showed that activation of FXR in L cells causes a reduction in glycolytic ATP production. They furthermore suggested that a decrease in transcription of multiple glycolytic enzymes in L cells were the cause of the reduced ATP production. In this study, we argue that FXR is not causing a decrease in transcription of multiple glycolytic enzymes. Instead, we find that FXR is causing the reduced glycolytic ATP production by targeting the glycolytic enzyme PKLR exclusively.

In an untargeted manner, we first identified the protein complex “merged-789” as significantly co-expressed, i.e. active, in FXR-activated L cells and control. The complex consisted of glycolytic enzymes, and we furthermore showed how the glycolysis as a whole was co-expressed as well. These findings indicated that the glycolysis in FXR-activated L cells was well-functioning and active; an opposite observation of what Trabelsi et al. had seen. Considering this controversy, we performed a DE analysis between control and FXR-activated L cells to test for significant downregulation of glycolytic enzymes. We used the expression data made publicly available by Trabelsi et al. and found no significant decrease in transcription of enzymes related to the glycolysis. Collectively, the glycolysis displays significant co-expression and show no differential expression, suggesting that the glycolysis is predominantly unaffected by the activation of FXR.

However, the observation of reduced glycolytic ATP production in L cells with activated FXR remains unchanged ^11^. Since FXR is not decreasing the expression of the glycolysis as a whole, another mechanism must exist. Looking for this mechanism, we went through glycolytic enzymes that showed deviating expression patterns. In this process, we found PKLR to stand out. PKLR, as the only essential enzyme, showed a decrease in co-expression with the other glycolytic enzymes in FXR-activated L cells relative to control. A decrease in co-expression suggests a decrease in activity ^19–23^. Furthermore, due to PKLR’s position in the glycolysis as the final enzyme, it may act as a bottleneck, reducing glycolytic activity altogether. We therefore hypothesised that PKLR is an indirect or direct target of FXR.

Turning to the literature, we looked for other studies mentioning FXR and PKLR. Interestingly, a study by *Caron et al*. in human hepatocytes studied PKLR during active FXR, and specifically found PKLR unresponsive to increasing glucose concentration. They suggest that FXR inhibits PKLR expression by blocking the transcription factor ChREBP. Despite the difference in cell type and organism, FXR’s mode-of-action is likely the same, as protein function in general is highly conserved ^30,31^. Our results and the *Caron et al*. findings can thus be seen as supportive evidence. We therefore find that PKLR is the most likely target of active FXR given the evidence available, and we suggest it causes the reduction in glycolytic ATP production (**Figure 4D**).

The glycolysis is deeply important in L cell biology, as the secretion of the insulin-stimulating hormone, GLP-1, is dependent on glycolytic activity ^32^. Specifically, an increase in glycolytic activity causes an increase in GLP-1 secretion ^32^. Describing how the glycolysis is repressed, in this case by FXR, is therefore an important part of delineating the key glycolytic enzymes linked to GLP-1 secretion. Growing interest in applying GLP-1 as pharmacotherapy emphasizes this importance. For T2D, development of GLP-1 receptor agonists has been a common choice of therapeutic strategy ^7,10^. However, other strategies have been suggested as well ^11^. Building on our results, we hypothesize that by protecting PKLR expression from the repressive effect of active FXR, decreased secretion of GLP-1 will not take place. Over time, this will increase GLP-1 secretion, as demonstrated before ^11^. Furthermore, protection of PKLR expression may be accomplished by preventing the binding of FXR to the transcription factor ChREBP. ChREBP have been shown to stimulate PKLR expression in human hepatocytes ^29^, which it cannot do when FXR is bound. Disruption of this binding will allow ChREBP to stimulate PKLR expression and consequently “protect” PKLR from FXR’s repressive effect.

## Author contributions

The study was a combined effort by KN and SB.

## Acknowledgements

Associate professor Kirstine G. Belling of the Novo Nordisk Foundation Center for Protein Research, University of Copenhagen, provided proof reading and helped structuring the paper.

## References

1. Reimann, F. et al. Glucose Sensing in L Cells: A Primary Cell Study. Cell Metab. 8, 532–539 (2008).

2. Gribble, F. M. & Reimann, F. Enteroendocrine Cells: Chemosensors in the Intestinal Epithelium. Annu. Rev. Physiol. 78, 277–299 (2016).

3. Grosse, J. et al. Insulin-like peptide 5 is an orexigenic gastrointestinal hormone. PNAS 111, 11133–11138 (2014).

4. Thomas, C. et al. TGR5-Mediated Bile Acid Sensing Controls Glucose Homeostasis. Cell Metab. 10, 167–177 (2009).

5. Hirasawa, A. et al. Free fatty acids regulate gut incretin glucagon-like peptide-1 secretion through GPR120. Nat. Med. 11, 90–94 (2005).

6. Greiner, T. U. & Bäckhed, F. Microbial regulation of GLP-1 and L-cell biology. Mol. Metab. 5, 753–758 (2016).

7. Meier, J. J. GLP-1 receptor agonists for individualized treatment of type 2 diabetes mellitus. Nat. Rev. Endocrinol. 8, 728–742 (2012).

8. Holst, J. J. The Physiology of Glucagon-like Peptide 1. Physiol. Rev. 87, 1409–1439 (2007).

9. Nauck, M., Stockmann, F., Ebert, R. & Creutzfeldt, W. Reduced incretin effect in Type 2 (non-insulin-dependent) diabetes. Diabetologia 29, 46–52 (1986).

10. Nauck, M. A., Vardarli, I., Deacon, C. F., Holst, J. J. & Meier, J. J. Secretion of glucagon-like peptide-1 (GLP-1) in type 2 diabetes: What is up, what is down? Diabetologia 54, 10–18 (2011).

11. Trabelsi, M.-S. et al. Farnesoid X receptor inhibits glucagon-like peptide-1 production by enteroendocrine L cells. Nat. Commun. 6, 7629 (2015).

12. Ananthanarayanan, M., Balasubramanian, N., Makishima, M., Mangelsdorf, D. J. & Suchy, F. J. Human Bile Salt Export Pump Promoter Is Transactivated by the Farnesoid X Receptor / Bile Acid Receptor. J. Biol. Chem. 276, 28857–28865 (2001).

13. Drucker, D. J., Jin, T., Asa, S. L., Young, T. A. & Brubaker, P. L. Activation of Proglucagon Gene Transcription by Protein Kinase-A a Novel Mouse Enteroendocrine Cell Line. Mol. Endocrinol. 8, 1646–1655 (1994).

14. Barabási, A-L.. & Oltvai, Z. N. Network biology: understanding the cell’s functional organization. Nat. Rev. Genet. 5, 101–113 (2004).

15. Kustatscher, G., Grabowski, P. & Rappsilber, J. Pervasive coexpression of spatially proximal genes is buffered at the protein level. Mol. Syst. Biol. 13, 937 (2017).

16. Li, T. et al. A scored human protein-protein interaction network to catalyze genomic interpretation. Nat. Methods 14, 61–64 (2017).

17. Li, H. & Liang, S. Local network topology in human protein interaction data predicts functional association. PLoS One 4, (2009).

18. Nepusz, T., Yu, H. & Paccanaro, A. Detecting overlapping protein complexes in protein-protein interaction networks. Nat. Methods 9, 471–472 (2012).

19. Pedersen, H. K., Gudmundsdottir, V. & Brunak, S. Pancreatic Islet Protein Complexes and Their Dysregulation in Type 2 Diabetes. Front. Genet. 8, 1–16 (2017).

20. Taylor, I. W. et al. Dynamic modularity in protein interaction networks predicts breast cancer outcome. Nat. Biotechnol. 27, 199–204 (2009).

21. Börnigen, D. et al. Concordance of gene expression in human protein complexes reveals tissue specificity and pathology. Nucleic Acids Res. 41, e171 (2013).

22. Kirk, I. K. et al. Chromosome-wise Protein Interaction Patterns and Their Impact on Functional Implications of Large-Scale Genomic Aberrations. Cell Syst. 4, 1–8 (2017).

23. Lee, S. et al. Network analyses identify liver-specific targets for treating liver diseases. Mol. Syst. Biol. 13, 938 (2017).

24. Ritchie, M. E. et al. limma powers differential expression analyses for RNA-sequencing and microarray studies. Nucleic Acids Res. 43, e47 (2015).

25. Reimand, J. et al. g:Profiler—a web server for functional interpretation of gene lists (2016 update). Nucleic Acids Res. 44, W83–W89 (2016).

26. Han, J.-D. J. et al. Evidence for dynamically organized modularity in the yeast protein-protein interaction network. Nature 430, 88–93 (2004).

27. Sartor, M. a et al. Intensity-based hierarchical Bayes method improves testing for differentially expressed genes in microarray experiments. BMC Bioinformatics 7, 538 (2006).

28. Hagberg, A. A., Schult, D. A. & Swart, P. J. Exploring Network Structure, Dynamics, and Function using NetworkX. Proc. 7th Python Sci. Conf. 1, 11–15 (2008).

29. Caron, S. et al. Farnesoid X Receptor Inhibits the Transcriptional Activity of Carbohydrate Response Element Binding Protein in Human Hepatocytes. Mol. Cell. Biol. 33, 2202–2211 (2013).

30. Dolinski, K. & Botstein, D. Orthology and Functional Conservation in Eukaryotes. Annu. Rev. Genet. 41, 465–507 (2007).

31. Shay, T. et al. Conservation and divergence in the transcriptional programs of the human and mouse immune systems. PNAS 110, 2946–2951 (2013).

32. Reimann, F. et al. Glucose Sensing in L Cells: A Primary Cell Study. Cell Metab. 8, 532–539 (2008).

